# Characterizing neuroinvasion and neuropathology of SARS-CoV-2 by using AC70 human ACE2 transgenic mice

**DOI:** 10.1101/2024.06.21.600068

**Authors:** Jason C. Hsu, Panatda Saenkham-Huntsinger, Pinghan Huang, Cassio Pontes Octaviani, Aleksandra K. Drelich, Bi-Hung Peng, Chien-Te K. Tseng

## Abstract

COVID-19 presents with a plethora of neurological signs and symptoms despite being characterized as a respiratory disease, including seizures, anxiety, depression, amnesia, attention deficits, and alterations in consciousness. The olfactory nerve is widely accepted as the neuroinvasive route by which the etiological agent SARS-CoV-2 enters the brain, but the trigeminal nerve is an often-overlooked additional route. Based on this consensus, we initially conducted a pilot experiment investigating the olfactory nerve route of SARS-CoV-2 neuroinvasion via intranasal inoculation in AC70 human ACE2 transgenic mice. Notably, we found that the trigeminal ganglion is an early and highly efficient site of viral replication, which then rapidly spread widely throughout the brain where neurons were primarily targeted. Despite the extensive viral infection across the brain, obvious evidence of tissue pathology including inflammatory infiltration, glial activation, and apoptotic cell deaths were not consistently observed, albeit inflammatory cytokines were significantly induced. However, the expression levels of different genes related to neuronal function, including the neurotransmitter dopamine pathway as well as synaptic function, and markers of neuronal damage were altered as compared to mock-infected mice. Our findings suggest that the trigeminal nerve can be a neuroinvasive route complementary to the olfactory nerve and that the ensuing neuroinvasion presented a unique neuropathological profile. This study provides insights into potential neuropathogenic mechanisms utilized by coronaviruses.

**IMPORTANCE:** COVID-19 presents with extrapulmonary signs and symptoms, the most notable of which involve the central nervous system, such as seizures and alterations in consciousness, and can eventually lead to death if severe enough. Some neurological signs and symptoms may continue to persist in some patients even after the resolution of active viral infection in the form of post-acute sequelae. Since the trigeminal nerve is a commonly under-studied route of entry into the brain in studies of coronaviruses and the neuropathogenic mechanisms of COVID-19 are not entirely elucidated, there is a need to thoroughly investigate this route of neuroinvasion. The significance of our research is in providing insights into the possible routes of SARS-CoV-2 neuroinvasion as well as the discovery of potential neuropathogenic mechanisms which may help guide the development of novel medical countermeasures.

## INTRODUCTION

Over four years now since its emergence in Wuhan, China(1) Coronavirus Infectious Disease-2019 (COVID-19) has infected over 700 million people in total across the globe, and killed over 6.9 million of them(2). The etiological agent of COVID-19 is a member of the coronavirus family in the *Betacoronavirus* genus designated ‘SARS-CoV-2.’(3) Despite being characterized as a primarily respiratory disease, COVID-19 importantly also exhibits neurological symptoms that range from the mild, such as loss of sense of smell (anosmia) and taste (ageusia) as well as headache and fatigue, to the severe, such as strokes and seizures, as well as neuropsychiatric disorders including delirium, anxiety, depression, psychosis, memory loss (amnesia), and attention deficits(4–12); with the plethora of neurological symptoms, it is undeniable that there is neurological system involvement in the pathogenesis of COVID-19.

After having caused much death and illness globally, COVID-19 continues to be a public health issue in the form of lingering neurological symptoms and post-acute sequelae(^13, 14^). Yet, despite great strides in the research effort in recent years, many of the neuropathogenic mechanisms of COVID-19 continue to elude elucidation, which would help in informing treatment of the neurological disorders that arise after disease onset.

One such neuropathogenic mechanism that still merits investigation is the routes of neuroinvasion taken by SARS-CoV-2 (hereby abbreviated to ‘SARS-CoV-2’). There is plentiful evidence to suggest SARS-CoV-2 has significant neuroinvasive potential(^4–6, 10–12, 15–18^). SARS-CoV-2 viral proteins and RNA have been detected within the brains of both COVID-19 patients and SARS-CoV-2-infected mice, leading to associated tissue pathology, such as microglial activation and immune infiltrates(^10, 15, 19, 20^). Furthermore, neural tissue cells, including neurons and glia, express low but sufficiently detectable levels of angiotensin-II-converting enzyme (ACE2) as well as transmembrane serine protease 2 (TMPRSS2), which have been identified as the main host enzymatic cofactors determining viral entry into permissive host cells(21–23).

Although at least eight routes of neuroinvasion have been hypothesized to be utilized by SARS-CoV-2(^4–6, 10–12, 15–18^), the current scientific consensus is the direct olfactory nerve is the main route of SARS-CoV-2 neuroinvasion, owing to the immediate exposure of the nerve endings to the outside atmosphere and short lengths of the nerves leading to the close proximity of the olfactory bulb (OB) to the external environment(19). Nevertheless, an oft-understudied route of neuroinvasion that should be considered here in the context of SARS-CoV-2 infection is the trigeminal nerve. Most well-characterized in studies involving human herpesvirus infections, particularly serotypes 1/2/3/6 (HHV-1/2/3/6)(24–27), the trigeminal nerve route of neuroinvasion is a particularly attractive route of neuroinvasion because the trigeminal neurons are directly connected to the brainstem at the pons while the nerve terminals still end in very close proximity to the external environment, only being separated by nasal epithelial cells, which have been reported to be susceptible to infection by SARS-CoV-2, thereby bypassing the blood-brain barrier (BBB)(^19, 28–31^).

There have been a few studies that seemingly confirm the trigeminal nerve route of neuroinvasion by SARS-CoV-2. Early in the pandemic, a study reported the detection of SARS-CoV-2 viral genome copies within the human TG while also reporting the detection of viral proteins in the human olfactory epithelium (OE) and OB(19). Similarly, a different study in K18 human ACE2 (hACE2) transgenic mice also reported the detection of SARS-CoV-2 viral genomes and infectious virions within the TG and brains, thus apparently validating the trigeminal nerve route of viral transmission(15). Based on these early reports, we conducted a pilot experiment investigating the olfactory nerve route of neuroinvasion to determine the regions of greatest viral tropism within the brains of hACE2 transgenic mice; it was during this pilot experiment that we incidentally found that the TG was intensely infected as well. In more recent studies, TG infection by SARS-CoV-2 has also been reported in deer mice(32). However, these previous studies failed to thoroughly investigate the trigeminal nerve route of neuroinvasion by SARS-CoV-2 and illustrate the implications of this route of SARS-CoV-2 neuroinvasion on COVID-19 pathogenesis in great detail. Based on these early findings and our observations in the preliminary exploratory experiment, we present here our thorough and detailed findings confirming the trigeminal nerve route as an early and efficient route of SARS-CoV-2 neuroinvasion in hACE2 transgenic mice, thereby resulting in a highly neurovirulent and neurotropic viral infection that induces altered neural function without obvious neuroinflammation or cell death.

## RESULTS

### Intranasal challenge with a lethal dose of SARS-CoV-2 caused a profound infection in the TG before spreading to the brain

To gain insights into the neuroinvasive potential of SARS-CoV-2, we intranasally challenged AC70 human ACE2 (hACE2) transgenic mice with 1×10^3^ TCID_50_ (approximately 333 LD_50_) of SARS-CoV-2 (US-WA-1/2020 strain) (Drelich et al., 2024; manuscript in press) and monitored them daily for morbidity (e.g., weight changes) and mortality. Starting on 4 days post-infection (dpi), the challenged mice began to exhibit significant weight loss as well as other signs of disease, before rapidly succumbing to infection with nearly 100% mortality by 5 dpi (Fig. 1A-C). Then, we assessed the kinetics of viral spread within the brain and its nearby peripheral nervous structures, e.g. OE and TG, by using immunohistochemical (IHC) staining for SARS-CoV-2 Spike (S) protein.

**Figure 1.**
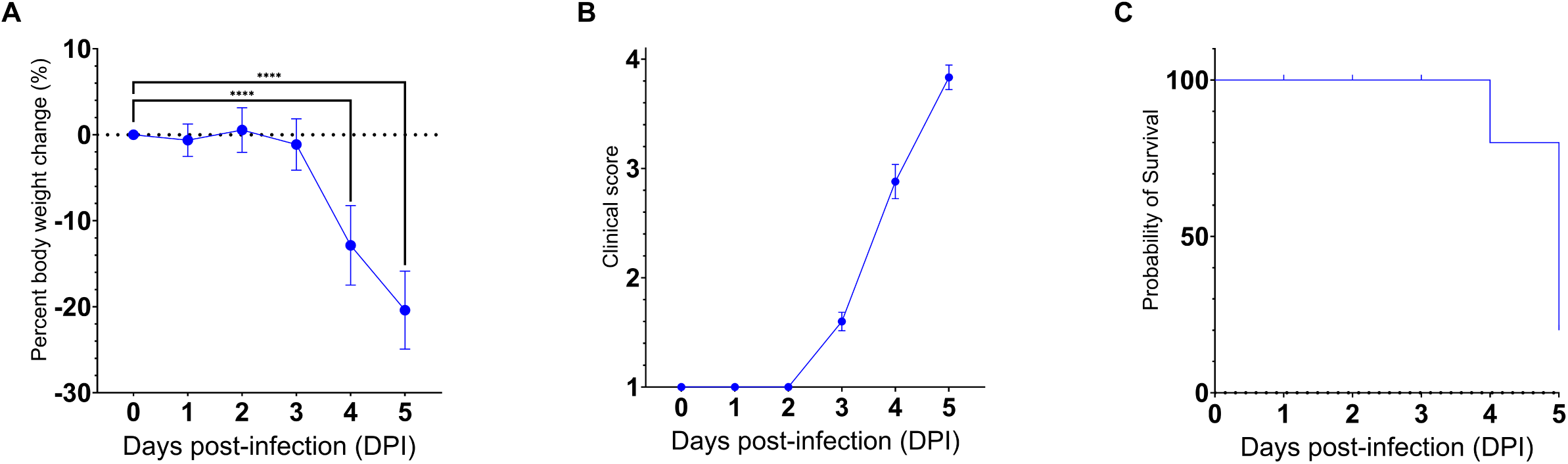
SARS-CoV-2 infection via the intranasal route results in a rapid development of clinical disease and mortality. 25 AC70 mice were intranasally inoculated with 1×10^3^ TCID_50_ of SARS-CoV-2 strain US-WA-1/2020 in 2% fetal bovine serum-supplemented cell media (2-MEM) and then five mice were sacrificed each day for five days post-infection. (A) Significant weight loss rapidly developed starting on 4 dpi, correlating with (B) the onset of severe clinical signs of disease. (C) The onset of clinical disease quickly advanced to near full mortality at 5 dpi Weight changes were expressed as the mean percent changes in infected animals relative to the initial weights at 0 dpi Error bars represent standard errors of the mean (SEM). ****p < 0.0001. These were the representative results of one out of two independent experiments.

We noted that SARS-CoV-2 S could only be sporadically detected within the OE as early as 1 dpi (one out of five mice), and progressively sustained thereafter through 4 dpi (Fig. 2A-D), thereby confirming earlier reports that the OE serves as an early site of viral replication(^19, 33^).

**Figure 2.**
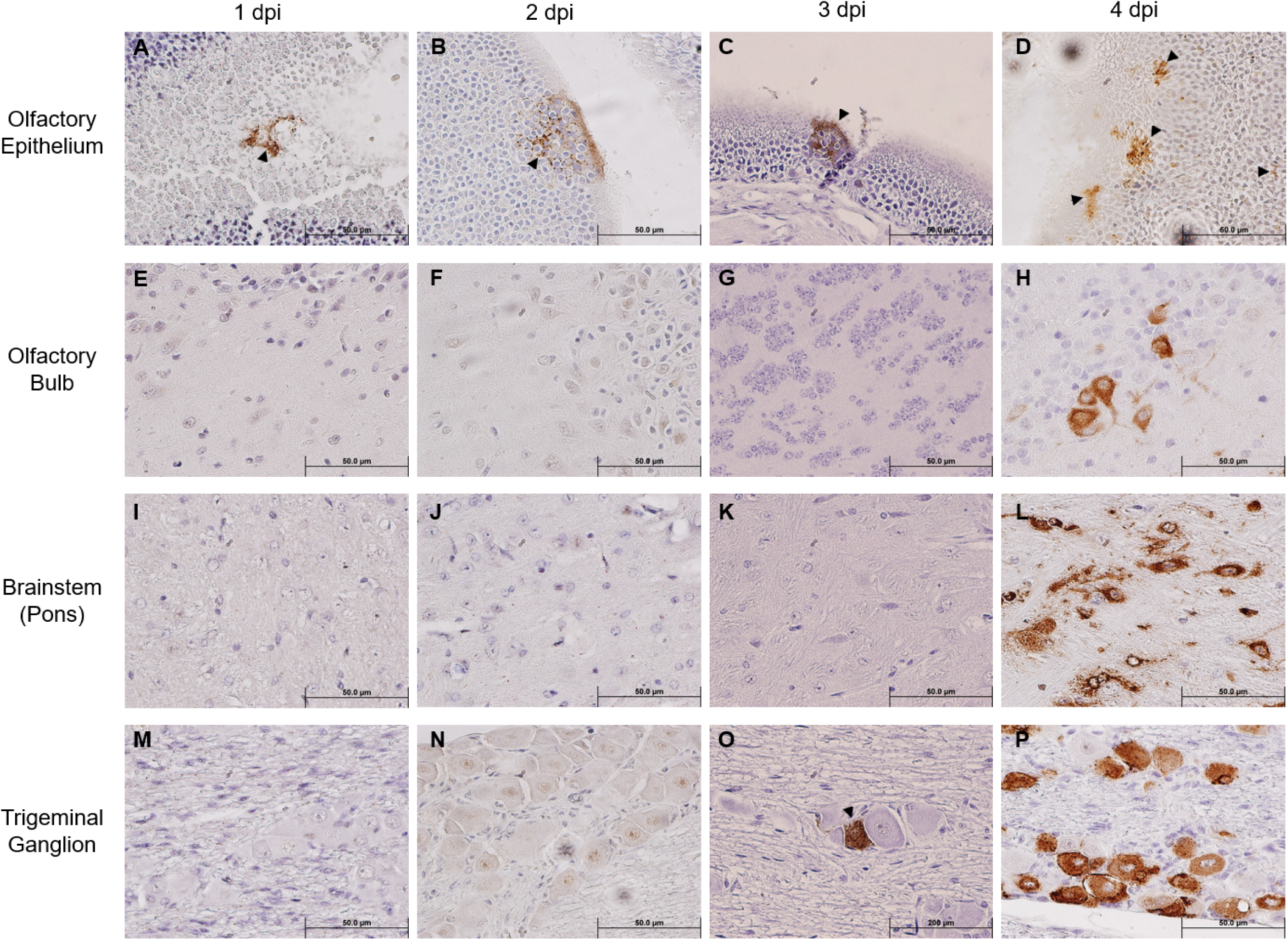
Immunohistochemical analysis of SARS-2 antigen in the brain, OE, and TG after infection via the intranasal route. Formalin-fixed, paraffin-embedded (FFPE) skull sagittal or coronal sections containing the brain, OE, and TG were analyzed via immunohistochemistry (IHC) for the expression of the SARS-2 spike (S) protein. SARS-2 S antigen (brown) can be detected only in the OE starting from 1 dpi but can begin to be detected in the TG starting from 3 dpi; SARS-2 S could not be detected in all other regions of the brain from 1 dpi to 3 dpi, including the olfactory bulb. Black arrowheads indicate selected points of antigen detection. (A-D) Olfactory epithelium; (E-H) olfactory bulb; (I-L) brainstem (pons); (M-P) trigeminal ganglion. Magnification 40X. Blue nuclei indicate hematoxylin counter-staining.

Despite the incrementing viral infection of OE over time, we were unable to detect any SARS-CoV-2 S within the OB until 4 dpi (Fig. 2E-H). Similarly, we could not detect any SARS-CoV-2 S staining within the pons either until 4 dpi (Fig. 2I-L). An analysis revealed that at 4 dpi most challenged mice stained positively for SARS-CoV-2 S in the OB and the pons. Interestingly, we were able to unambiguously detect the expression of SARS-CoV-2 S in the TG starting at 3 dpi in a couple of infected mice, of which the staining intensity profoundly increased in all infected mice at 4 dpi (Fig. 2M-P). Taken together, the finding of TG as a permissive site of SARS-CoV-2 infection suggests that the trigeminal nerve route could be another early and olfactory nerve-independent route of neuroinvasion by SARS-CoV-2 to enter the CNS/brain. Nevertheless, consistent with the onset of severe disease in infected mice, we observed at 4 dpi overwhelming viral infection in all major anatomic regions of the brain, including the proposed initial ports of entry, i.e., OB and pons of the olfactory nerve and trigeminal nerve neuroinvasive routes, respectively (Fig. 2H&L), as well as regions distal to the initial sites of entry, such as the frontal cerebral cortex (prefrontal, somatomotor, somatosensory, etc.), basal ganglia (caudate putamen and striatum), thalamus, hypothalamus, hippocampal formation, cerebellum, and brainstem (mesencephalon and medulla) (Suppl. Fig. 2). At 5 dpi, we observed the viral antigen staining continue to spread throughout almost all regions of the brain (Suppl. Fig. 3).

### Kinetics of SARS-CoV-2 viral infection in brain and TG

As we have shown the TG is an early site of SARS-CoV-2 infection, we investigated the kinetics of viral infection within the brain and TG. As shown in Figure 3, we found that infectious virus could be recovered from the brain at 3 dpi with a titer of approximately 4.5 log TCID_50_/g, followed by a sharp increase to ∼7 log and ∼7.5 log TCID_50_/g at 4 and 5 dpi, respectively. While we could not detect any signs of viral infection by IHC staining within the TG until 3 dpi (Fig. 2O), infectious virus was recovered at 2 dpi (∼4 log TCID_50_/g). The titers of infectious virus in the TG increased thereafter to 6.5 log TCID_50_/g 5 dpi. Specifically, while we were only able to isolate a low titer of live virus from the TG of 1/10 challenged mice at 1 dpi, we were able to increase the detection frequence to 7/10 and 10/10 at 2 and 3 dpi and thereafter. Therefore, viral replication occurs one day earlier in the TG than in the brain.

**Figure 3.**
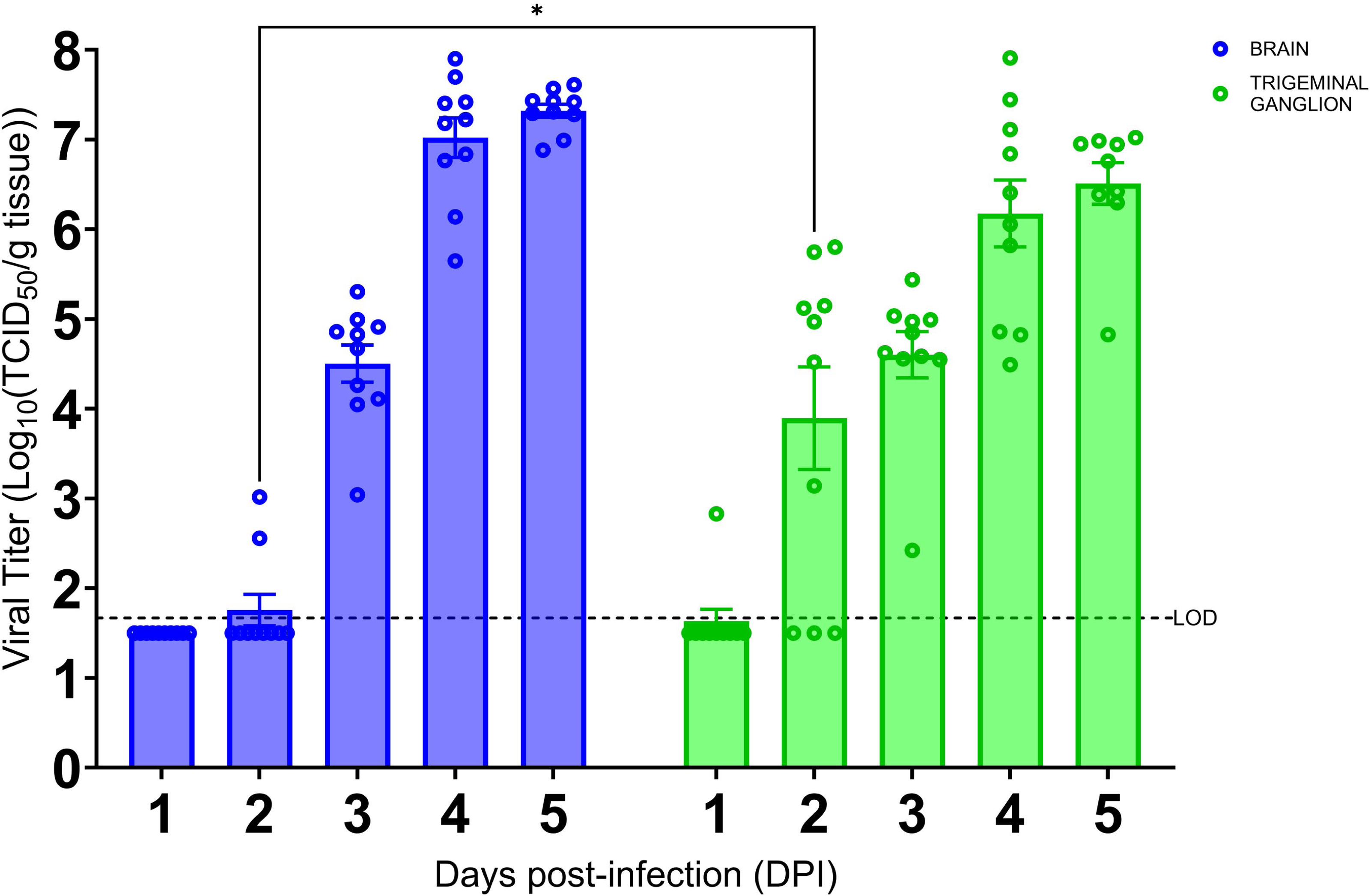
SARS-CoV-2 replication kinetics in the trigeminal ganglion and the brain. The titers of infectious virus in brain and TG were calculated and expressed as log_10_ TCID_50_ virus per gram of tissue and were plotted as the mean of two different cohorts (n = 10 animals per timepoint). Virus titers in the brain (blue) and TG (green) were assessed using a standard Vero-E6 cell-based TCID_50_ assay. *p < 0.05, by Student’s *t*-test, comparing brain and TG. Error bars represent standard errors of the mean (SEM). These were the combined data of two different independent experiments.

### Neurons are the primary brain cells supporting productive SARS-CoV-2 infection

The profound SARS-CoV-2 infection within the brain of infected mice prompted us to investigate the identity of permissive brain cells by using two-color immunofluorescent (IF) staining by simultaneously targeting specific cell markers and viral antigens. Encouraged by the data shown in Fig. 2 and Suppl. Fig. 3 that a vast majority of cells positively stained for the S protein by IHC morphologically resembled neuronal cells, we repeated the IHC staining for the expression of beta tubulin III (TUBB3, also known as Tuj1), a marker of neuronal cells, and SARS-CoV-2 S protein. We found that most cells that were stained positively for the S protein co-expressed Tuj1 in the cytosols of the main bodies of the neurons, indicating neurons are likely the prime brain cells permissive to SARS-CoV-2 infection (Fig. 4A-D). To further verify that neurons were indeed the preferred brain cells targeted by SARS-CoV-2, we use the same IHC staining technique for detecting the expressions of glial fibrillary acidic protein (GFAP) and ionized calcium-binding adaptor molecule 1 (IBA1), markers for astrocytes and microglia, respectively, along with SARS-CoV-2 S protein. While there were a few cells co-labelled with GFAP and SARS-CoV-2 S (Fig. 4H, arrowheads), the majority of GFAP^+^ astroglia were not permissive to SARS-CoV-2 infection (Fig. 4E-H). Moreover, we did not observe any signs of proliferative response (astrogliosis) and activation of astroglia (Fig. 4F), based on the absence of detectable extension and thickening of cellular processes(34–36), when compared to mock-infected animals (Fig. 4J). In contrast to SARS-CoV-2-permissive neurons and, astroglia, to a much lesser extent, we were unable to reveal any cells dually labelled with IBA1, the marker of microglial cells, and SARS-CoV-2 S protein (Fig. 4M-P), indicating that microglial cells likely are not permissive to infection by SARS-CoV-2. However, as we could only detect very few cells that were IBA1^+^, we could not make any conclusions on microglial activation based on proliferation and retraction of processes (Fig. 4N).

**Figure 4.**
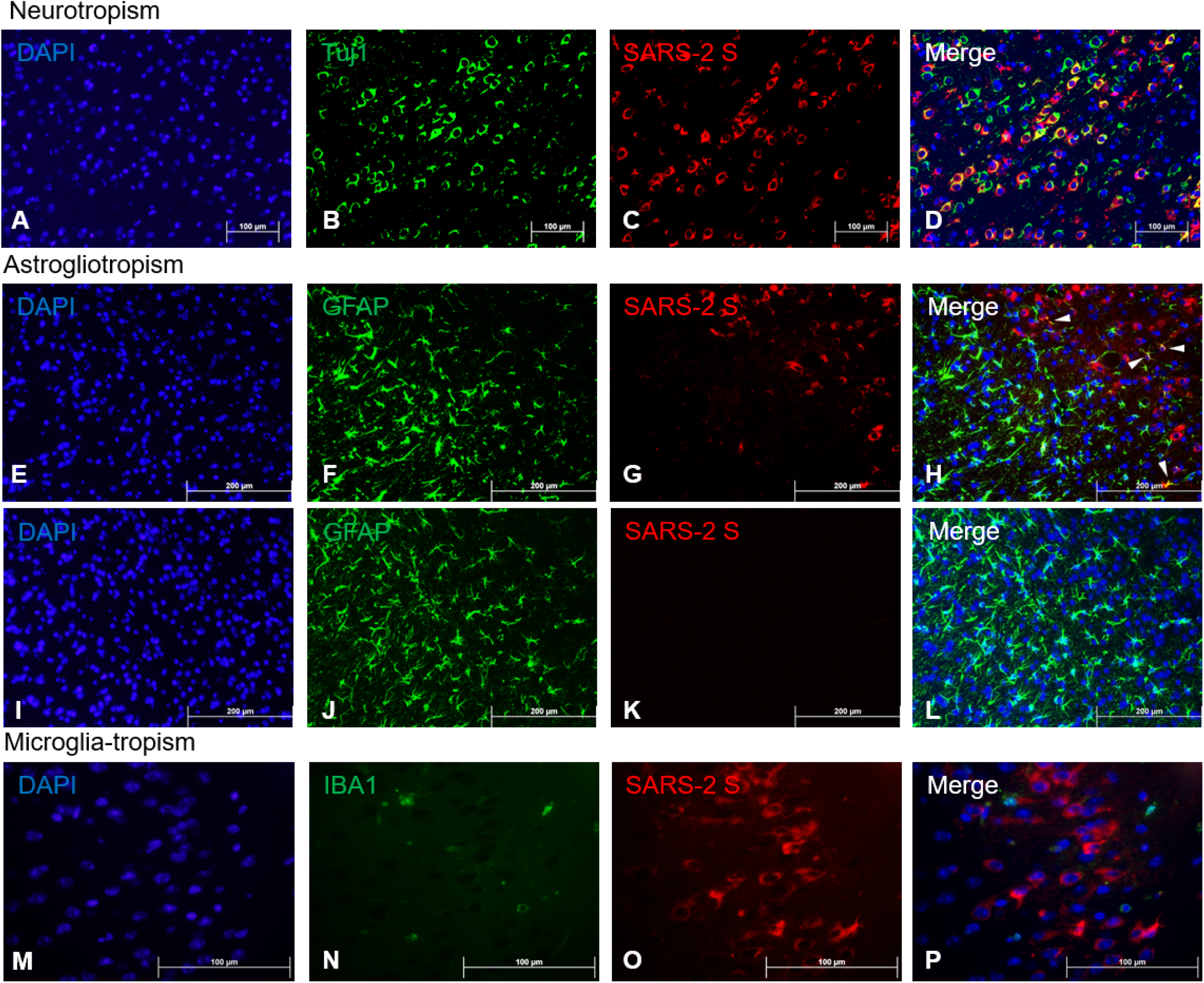
Viral tropism analysis via immunofluorescence of SARS-CoV-2 antigen in the brain. Serial sections of the FFPE brain sections were analyzed via dual-labeling immunofluorescence (IF) for the expression of the SARS-2 spike (S) protein (red) and different cell identity markers (green). (A-D) Neurons (Tuj1^+^, frontal cortex); (E-L) Astrocytes (GFAP^+^, frontal cortex), white arrowheads indicate selected points of colocalization, E-H: SARS-CoV-2-infected mice, I-L: mock-infected mice; (M-P) Microglia (IBA1^+^, frontal cortex). Magnifications: A to L, 10X; M to P, 40X. DAPI counterstaining (blue).

While human (h) ACE2 transgene is known to constitutively express in tissues/organs of AC70 transgenic mice(^37, 38^), to what extent this human hACE2 expression conferred the susceptibility of brain cells to SARS-CoV-2 infection has not been fully investigated. To study this, paraffin-embedded brain sections of infected AC70 mice were subjected to the standard IHC staining for hACE2 and SARS-CoV-2 S protein, as described above. As shown in Fig. 5A-D, we found that in the choroid plexus, hACE2 expression alone cannot act as the determinant for permissiveness to SARS-CoV-2 infection; the choroid plexus, which is composed of endothelial and glial ependymal cells, was shown to intensely express hACE2 (Fig. 5B), which is consistent with the earlier reports(23), and yet, cells within this region apparently were not stained positively with SARS-CoV-2 S protein (Fig. 5C&D). The nature of such a loose correlation between hACE2 expression and SARS-CoV-2-permissiveness within different regions of the brain warrants additional studies.

**Figure 5.**
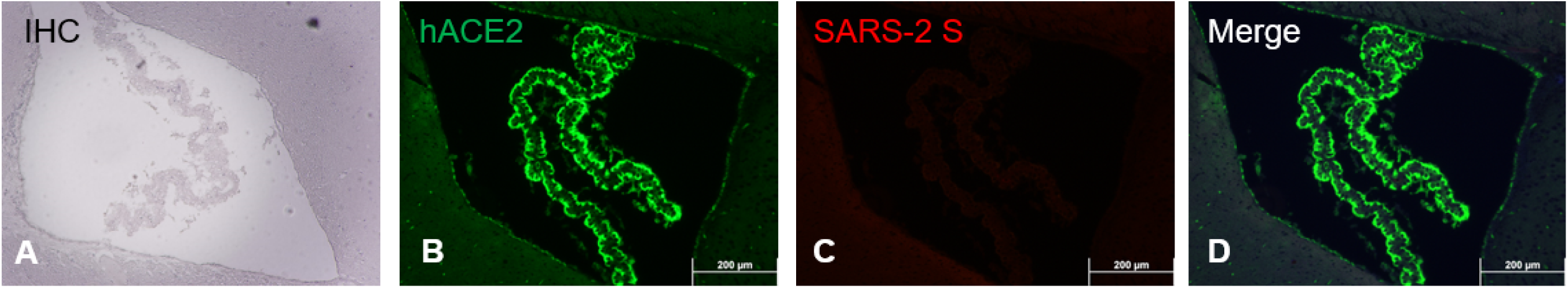
ACE2 co-expression with SARS-CoV-2 S antigen at the choroid plexus. FFPE brain section showing specifically the choroid plexus analyzed via immunostaining (IHC and dual-labeling IF) for hACE2 and SARS-2 S. (A) SARS-2 S IHC (brown); (B) hACE2 IF (ACE2^+^, green); (C) SARS-2 S IF (SARS-2 S^+^, red); (D) Merge of hACE2 and SARS-2 IF (ACE2^+^ and SARS-2 S^+^). Magnification 10X. IHC hematoxylin counterstaining (blue).

### Host responses to SARS-CoV-2 infection within the brains of AC70 transgenic mice

Having revealed the profound viral infection throughout major anatomical regions of the brain, we investigated how host would respond in the brain upon lethal challenge with SARS-CoV-2. We initially profiled the inflammatory responses by RT-qPCR, followed by examining the brain sections for the histopathology. We found that among 13 inflammatory mediators measured, 11 mediators were significantly induced in the brains at 4 dpi (Fig. 6), the time when significantly elevated viral titers were recovered, as shown in Fig. 3. We also found that 10 out of 11 virally induced soluble mediators were proinflammatory, including IFN-I (α/β), IFN-II (γ), TNF-α, IL-1β, IL-6, IP-10, MCP-1, MX-1, and RANTES, whereas the transcriptional levels of IL-4 and IL-10, markers of Th2 and anti-inflammatory or inflammatory regulator, respectively, were either slightly downregulated (IL-4) or significantly upregulated (IL-10). Despite the significant expression of inflammatory mediators within the brain, we did not identify any infiltrates of mononuclear cells within H&E-stained sections of the brains even at 4 dpi (Suppl. Fig. 1). Neither did we notice any readily detectable tissue damage in the brains. Because SARS-CoV-2 infection exhibits cytopathic effects resulting in the deaths of permissive host cells(39), we examined the brain sections for any histopathological signs of cell death. As shown in Suppl. Fig. 1, we were unable to detect any signs of apoptotic cell death as apoptotic cell deaths were the most common type of cell deaths associated with neurovirulent viral infections. We used the standard TUNEL assay kit (Abcam, Cambridge, UK) to detect apoptotic cells within the brains harvested at 5 dpi. Among a total of five brains examined, only one exhibited a few apoptotic cells while the other four were negative for TUNEL assays (data not shown). Together, these results suggested that infected brain cells, especially neurons as indicated by the IF dual-labeling results (Fig. 4A-D), but not inflammatory infiltrates, are the likely sources of the inflammatory mediators detected in the brain. Additionally, the lack of inflammatory infiltrates and convincing cell death within the brain emphasize the neuronopathy, but not encephalitis, is the likely cause of death of acutely infected AC70 transgenic mice.

**Figure 6.**
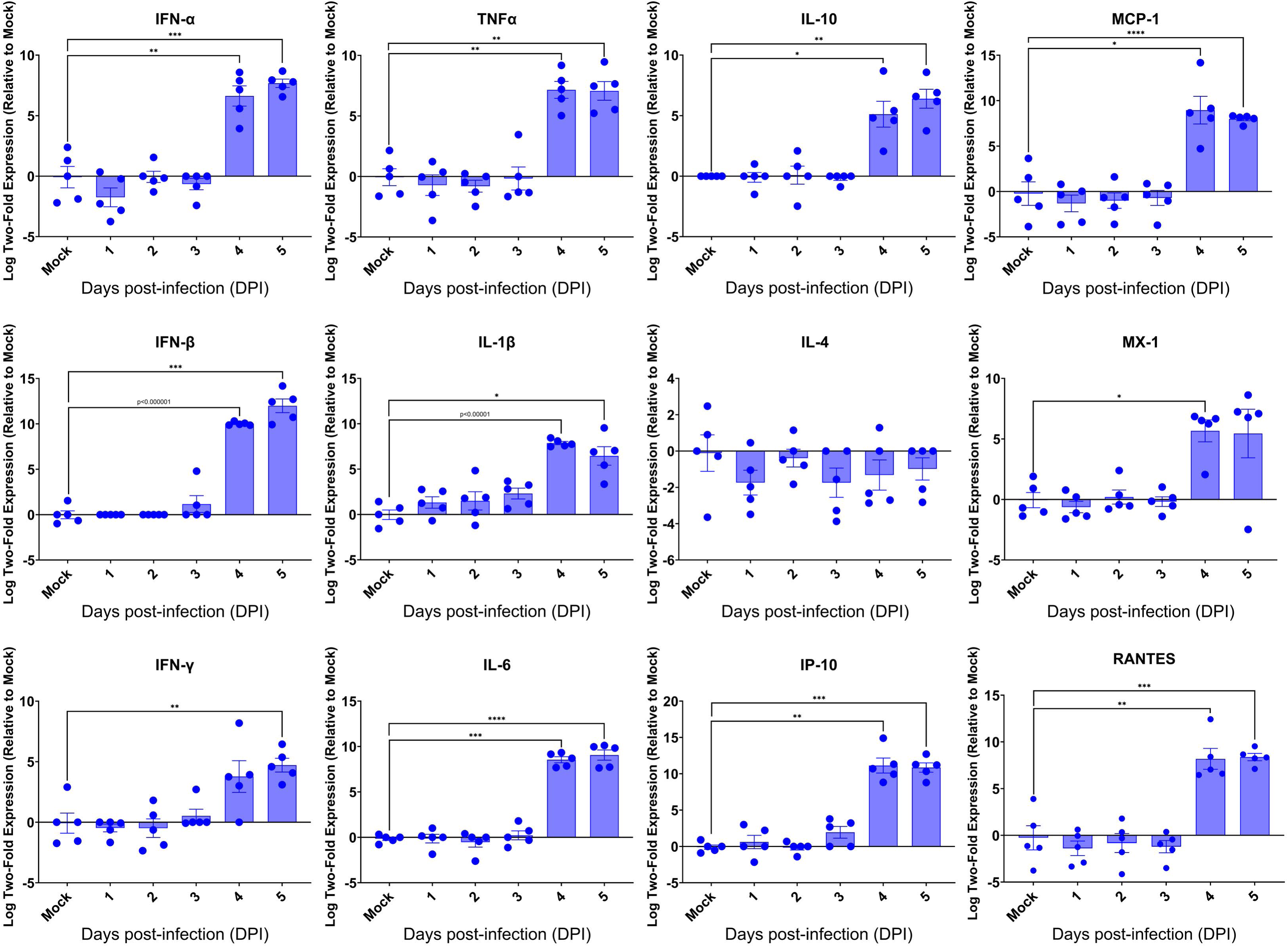
Kinetics of the cytokine responses in the brains of SARS-2-infected AC70 mice. Total RNA extracted from the brains of AC70 mice sacrificed daily after SARS-2 infection were used to measure the expression of various cytokines and chemokines by RT-qPCR. Each individual brain sample was assayed in duplicate. Results are shown as the mean for five animals at each time point. Error bars represent SEM. *p < 0.05, **p < 0.01, ***p < 0.001, ****p < 0.0001, p < 0.00001 where indicated (Student’s *t*-test, compared to mock-infected mice).

### Neuroinvasive SARS-CoV-2 infection dysregulated the expression of key genes regulating neurological functions

Despite the intense SARS-CoV-2 infection within the brain, preferentially targeting neurons, especially neurons, of acutely infected AC70 transgenic mice that succumbed to infection within days, we did not reveal histopathological evidence of neuroinflammatory response (encephalitis) with noticeable cell death. We examined whether this seemingly nonlytic, but extensive neuronal infection of SARS-CoV-2 might still alter neurological functions. Thus, we explored the expression of six key genes, i.e., *DRD1*, *TH*, *NEFL*, *Eno2* (*NSE)*, *Syn-1A*, *SNAP-25*, that serve as the regulators and markers for neuronal function and damage, respectively, in the brains of infected AC70 transgenic mice over time, compared to uninfected brains. We found that, except for *DRD1*, the transcriptional expressions of all other five genes evaluated were downregulated, with statistical significance only at a few timepoints for *TH*, *NEFL*, and *Syn-1A* (Fig. 8). The up- and down-regulated expressions of the dopamine receptor D1 (*DRD1*) and tyrosine hydroxylase (*TH*), respectively, a functional pair of molecules governing a unique neuronal function, is of interest; at 2 dpi, the significant upregulation of DRD1 was mirrored by the significant downregulation of TH (Fig. 7). Nevertheless, these results show that neuroinvasive SARS-CoV-2 infection could indeed alter functional gene expression without causing neuroinflammation or obvious cell death.

**Figure 7.**
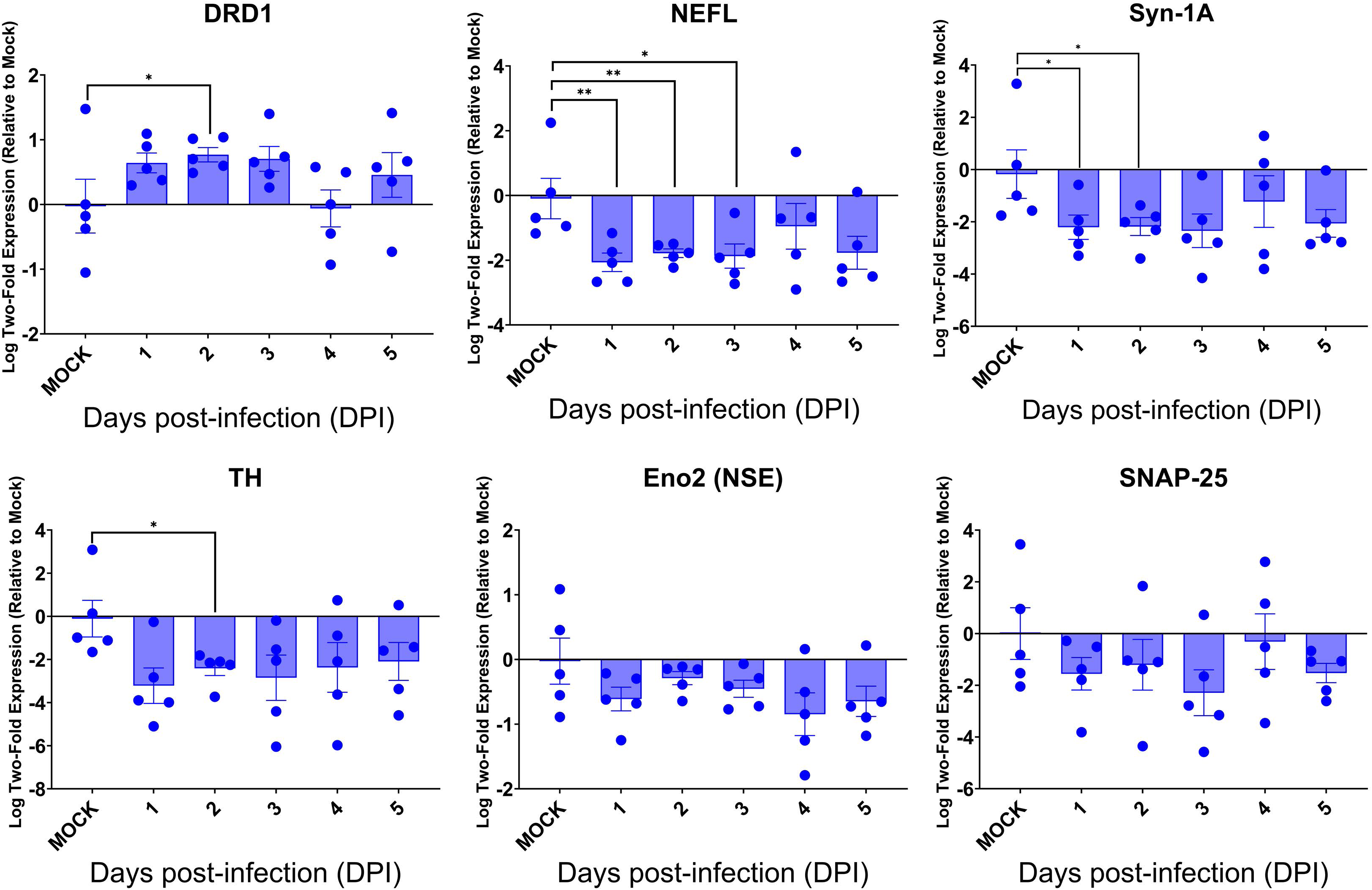
SARS-CoV-2 brain infection alters gene expression of neuronal function and neural damage markers. Total RNA extracted from the brains of infected AC70 mice daily after SARS-2 infection were used to measure the gene expression levels of selected neural biomarkers of damage (*Nefl* and *Eno2*) and neuronal function (*DRD1*, *TH*, *Syn-1A*, and *SNAP-25*). Each individual brain sample was assayed in duplicate. Results are shown as means (±SEM) of five animals at each time point. *p < 0.05, **p < 0.01 (Student’s *t*-test, compared to mock-infected mice).

**Figure 8.**
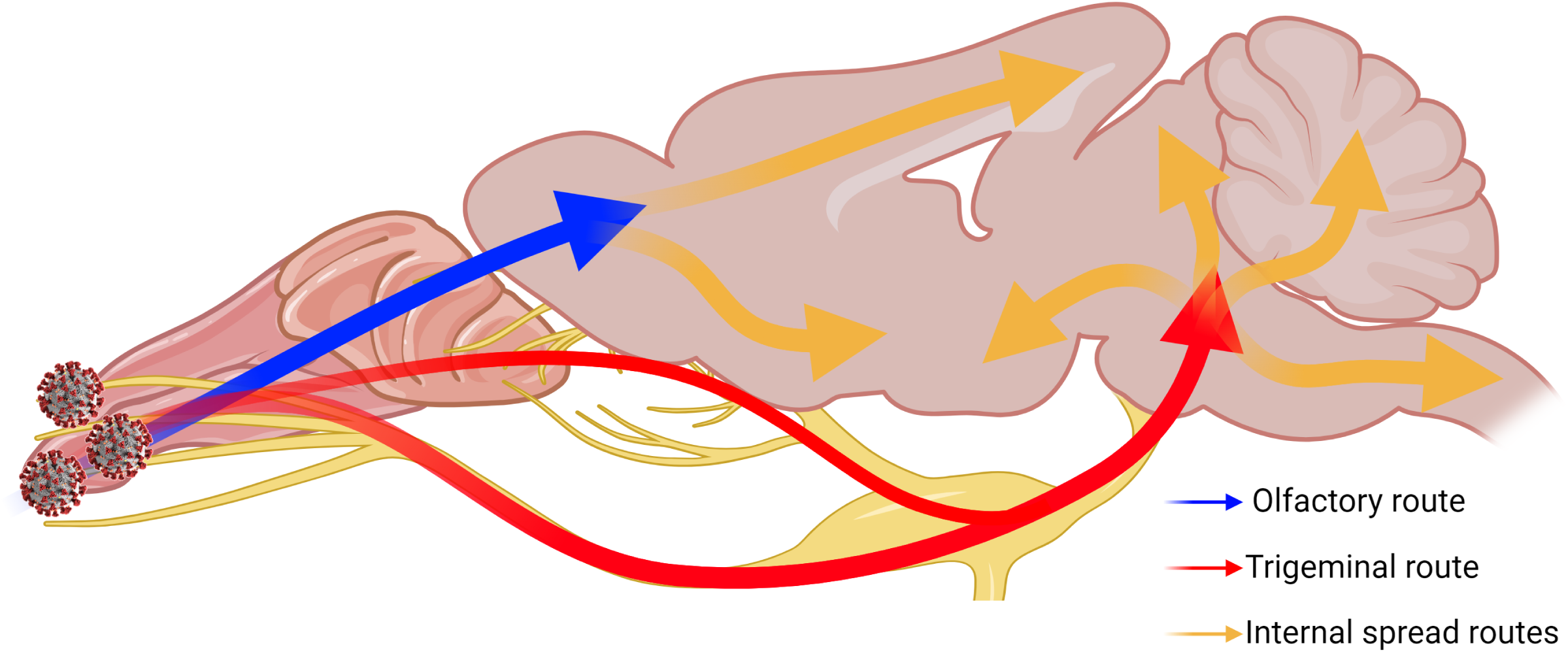
Two-pronged neuroinvasion route of SARS-CoV-2. Theorized two-pronged neuroinvasion route taken by SARS-CoV-2 starting from the nasal cavity during respiratory infection based on the observed viral antigen staining pattern. Red and blue arrows indicate the two main forks invading into the brain (e.g. OE → OB (blue); nasal/olfactory epithelium → TG → pons (red)). Orange arrows indicate speculated routes of further viral spread inside the brain proper once brain has been penetrated.

## DISCUSSION

In the study of coronavirus pathogenesis and particularly that of SARS-CoV-2, the trigeminal nerve is a possible route into the CNS that is often overlooked in favor of the olfactory nerve. Widely reported and generally accepted as the main route of entry into the CNS, the olfactory nerve route alone does not explain how SARS-CoV-2 infection could penetrate the main bulk of the brain at its rear(40–42). To this end, we mapped and characterized the trigeminal nerve route of neuroinvasion by SARS-CoV-2 in the AC70 hACE2 transgenic mouse model and found that there are many potential consequences. Specifically, we demonstrated that the trigeminal nerve may be an early and highly efficient site of SARS-CoV-2 viral replication, on par with that of the olfactory nerve, and that SARS-CoV-2 viral infection primarily targets neurons, leading to changes in neural function with minimal tissue pathology.

Early in the course of viral infection, SARS-CoV-2 can be readily detected in the TG, even before the onset of weight loss and disease signs in our mouse model. IHC staining showing the early SARS-CoV-2 S staining in the TG (Fig. 2O&P), supported by the high viral titer of the TG at an earlier timepoint (Fig. 3), implied that viral infection of the trigeminal nerve occurred nearly simultaneously with the OE and olfactory nerve, which would have occurred immediately after intranasal challenge. Additionally, the SARS-CoV-2 S staining pattern changing from undetectable to observable in all major anatomic regions of the brain within the span of one day suggested that the spread of viral infection occurred extremely rapidly. Since on 4 dpi the peripheral regions of the brain closest to both the OE (e.g. OB and frontal cortex) and the TG (e.g. hypothalamus, midbrain, and pons) (Suppl. Fig. 3A) stained more intensely with SARS-CoV-2 S than the central internal regions, we speculated that SARS-CoV-2 neuroinvasion proceeded via retrograde axonal transmission. Alternatively, the more extensive viral antigen staining of the peripheral anatomic regions of the brain than the central regions could indicate the virus was transported in the cerebrospinal fluid (CSF), as has been previously suggested(43); not only is the TG itself situated anatomically in the CSF-rich Meckel’s cavity, but terminals of all three branches of the trigeminal nerve penetrate through the cribriform plate to innervate skull bone marrow niches and ultimately the dura mater of the meninges, which are awash in CSF(^28, 44, 45^). Although the TG and trigeminal nerve were previously reported to have been infected early on during the heights of the COVID-19 pandemic(19), this study greatly extends the findings from the previous study by investigating the dynamics of SARS-CoV-2 neuroinvasion along the trigeminal nerve on a time course basis.

Our findings indicate that the primary target of SARS-CoV-2 viral tropism in the brain is neurons, but not astrocytes, microglia, or even endothelial cells. Initially, based on the expression levels of ACE2, in descending order, the main targets of SARS-CoV-2 tropism in the brain were thought to be endothelial cells, and then ependymal glial cells, astrocytes, and microglia, but not neurons(^46, 47^). Yet, our IF staining showing the amount of SARS-CoV-2 S expression in different cell types to be highest in neurons (Fig. 4A-D) suggests otherwise in our model. We identified an anatomic structure of the brain, the choroid plexus, as not only the structure of the brain with one of the highest expression levels of ACE2 as previously reported(^23, 48^), but one that was not observably infected by SARS-CoV-2 at all. We were interested in the permissiveness of the choroid plexus to SARS-CoV-2 because the choroid plexus has been suggested as an alternative portal of viral entry into the brain via the hematogenous routes due to its composition of primarily endothelial and ependymal glial cells(^10, 23, 48^).

Despite the profound viral infection throughout the brain, the brain was notably devoid of the typical correlates of tissue pathology associated with neuro-dysfunction. Although the pro-inflammatory cytokines were generally significantly induced corresponding to the magnitude and kinetics of viral replication, we were initially very surprised to observe an overall lack of any histopathological signs of neuroinflammation; inflammatory infiltrates and the associated vascular cuffing were rarely, if at all, observed, while our IF staining revealed a lack of astrocytic and microglial activation, which is atypical for viral neuroinvasion. However, we rationalized this by considering a sufficiently highly virulent and rapid viral infection has the ability to induce an immunosuppressive state that the neuroimmune system does not have the time to mount an inflammatory response before the organism succumbs(49–51). Additionally, we noticed that the occurrences of apoptosis were not consistent with the vast extent of viral infection; for example, in two anatomic regions of the brain with heavy viral infection, the frontal cortex and medulla oblongata, there were only a few cells actively undergoing apoptosis as revealed by TUNEL assay, while the hypothalamus, one of the most heavily infected region of the brain, had no cells detected undergoing apoptosis. However, we again surmised that owing to their postmitotic nature and limited numbers, mature neurons are remarkably resistant to apoptosis and programmed cell death, even after viral infections(52–54). All the histopathological results combined prompted us to investigate the expression levels of neural function markers, namely those for dopamine neurotransmitter processing and synaptic function, to gain any insights into the mechanisms of neurodysfunction. The pair of DRD1 and TH is relevant because this gene couple is directly involved in neurotransmitter dopamine processing; TH is the rate-limiting enzyme that catalyzes the production of the dopamine precursor L-DOPA(55), while DRD1 is the most abundant dopamine receptor within the CNS(56). Furthermore, the downregulation of the neural damage markers neurofilament light chain (*NEFL*) and neuron-specific enolase (*NSE*) (Fig. 7) ran counter to our expectations; in cases of acute brain injury, such as traumatic brain injuries (TBI) or ischemic events, NEFL and NSE have been reported to be elevated as markers of neuronal injury(57–61). These results indicated to us that there were indeed alterations to neural function because of viral infection, whether due to inflammation or viral infection. Our results suggest that there are alternative mechanisms of neurological disorder that extend beyond neuroinflammation and cell death.

Overall, our results show that the trigeminal nerve is an early and efficient site of SARS-CoV-2 infection in our model, suggesting that it may be an efficient entry route to the brain/CNS. Based on all our results together, we speculated neuroinvasive SARS-CoV-2 uses a two-pronged route from the OB and the pons towards the center of the brain (Fig. 8). We also characterized alterations in neuronal function that were observed despite a general lack of typical histopathological findings of neuroinflammation. It is clear from these findings that additional studies are warranted for the further characterization of COVID-19 neuropathogenesis.

## Materials and Methods

All procedures involving animals and infectious virus were performed in a biosafety level 3 (BSL-3) or animal biosafety level 3 (ABSL-3) facility at Galveston National Laboratory at the University of Texas Medical Branch (UTMB) at Galveston, Texas, an Association for Assessment and Accreditation of Laboratory Animal Care (AAALAC)-accredited (November 24, 2020) and Public Health Service-Office of Laboratory Animal Welfare (PHS-OLAW)-approved (February 26, 2021) high-containment National Laboratory. All animal procedures were carried out in accordance with animal protocols approved by an Institutional Animal Care and Use Committee (IACUC) at UTMB.

### Virus

The SARS-CoV-2 (strain US-WA-1/2020) used throughout this study was generously provided to us by Dr. Natalie Thornburg at the Centers for Disease Control (CDC), Atlanta, GA, through the World Reference Center for Emerging Viruses and Arboviruses (WRCEVA). SARS-CoV-2 were propagated in Eagle’s Minimal Essential Medium (MEM) (Corning, 10–010–CV) supplemented with 2% fetal bovine serum (FBS), 2% L-Glutamine (GIBCO, 25030–164), and 1% Penicillin-Streptomycin (GIBCO, 15140–122); this media formulation has been designated ‘2-MEM.’ The original stock of SARS-CoV-2 was cultured in 2-MEM and passaged two more times in Vero-E6 cells to generate the working viral stocks, which were stored at − 80 °C. The working viral stocks used throughout this study were titrated at ∼ 5 × 10^6^ TCID_50_/mL by a standard TCID_50_ assay in Vero-E6 cells.

### Cells

Vero-E6 immortalized African green monkey kidney cells (CRL-1580, American Type Culture Collection) were grown in a media formulation designated ’10-MEM,’ a media formulation similar to 2-MEM but supplemented instead with 10% FBS.

### SARS-CoV-2 infection and necropsy

Isoflurane-anesthetized female AC70 hACE2 transgenic mice at 8-9 weeks old were challenged intranasally with 1×10^3^ TCID_50_ SARS-CoV-2 in 60 µL of 2-MEM; five control mice were mock-challenged with the same volume of phosphate-buffered saline. All mice were weighed daily to monitor disease progression. Additionally, illness severity in infected mice was scored independently by two investigators who used a standardized 1-4 grading system as follows: 1, healthy; 2, ruffled fur, lethargic; 3, ruffled fur, lethargic, hunched posture, orbital tightening, labored breathing/dyspnea, and/or more than 15% weight loss; 4, reluctance to move when stimulated or at least 20% weight loss. Each day after infection, five infected mice were sacrificed to obtain whole skulls for determining viral infectivity titers, staining for viral antigen by IHC as well as other antigens by two-color IF, profiling inflammatory responses, and histopathological analysis. The control mice were sacrificed on the first day post-mock-challenge (1 dpi) to harvest the same as above described. The whole skull samples were then split into left and right hemispheres, with the left hemispheres subsequently being further split into whole brain and trigeminal ganglion samples; half of each of whole brain and trigeminal ganglion samples would later be homogenized in 2% FBS-PBS while the other half of each tissue type samples would be homogenized in TRIzol (Invitrogen, Waltham, Massachusetts, USA). The remaining right hemisphere of each whole skull sample was then fixed by immersion in 10%-buffered formalin for 72 hours followed by transfer to 70% ethanol.

### End-point dilution median tissue culture infectious dose (TCID_50_) viral titration assay

The end-point dilution median tissue culture infectious dose (TCID_50_) viral titration assay was performed as previously described(^37, 38, 62^). Briefly summarized, after an initial clarification by centrifugation step, we carried out a 1:10 serial dilution from 10^-1^ to 10^-8^ from a starting dilution of 50 µL of viral samples into 450 µL of 2-MEM. Then, we aliquoted 100 µL of the dilution into a 96-well plate of confluent Vero E6 cells at four wells each dilution. All 96-well plates were incubated at 37°C at 5% CO_2_ for up to three days, after which the number of wells exhibiting cytopathic effect were counted for each dilution. Then, the number of viable virions were calculated and quantified based on the Reed and Muench method and expressed as TCID_50_/mL(63).

### RNA extraction and reverse transcription-quantitative polymerase chain reaction (RT-qPCR)

Total RNA was isolated from the tissues of infected mice homogenized in TRIzol solution as indicated above using a chloroform extraction method according to manufacturer instructions. Contaminating genomic DNA was removed upon digestion with DNase I during the extraction procedure using a DNase I clean-up kit (Invitrogen, AM1907, Waltham, Massachusetts, USA). The resulting RNA samples were subjected to two-step RT-qPCR analysis to assess the expression of SARS-CoV-2 E gene as well as other genes, starting with reverse transcription into cDNA using the iScript Reverse Transcription kit (Bio-Rad, 1708841Hercules, California, USA). The primers for all genes can be seen in Table 1. 18S rRNA was used as the endogenous control.

**Table 1.**
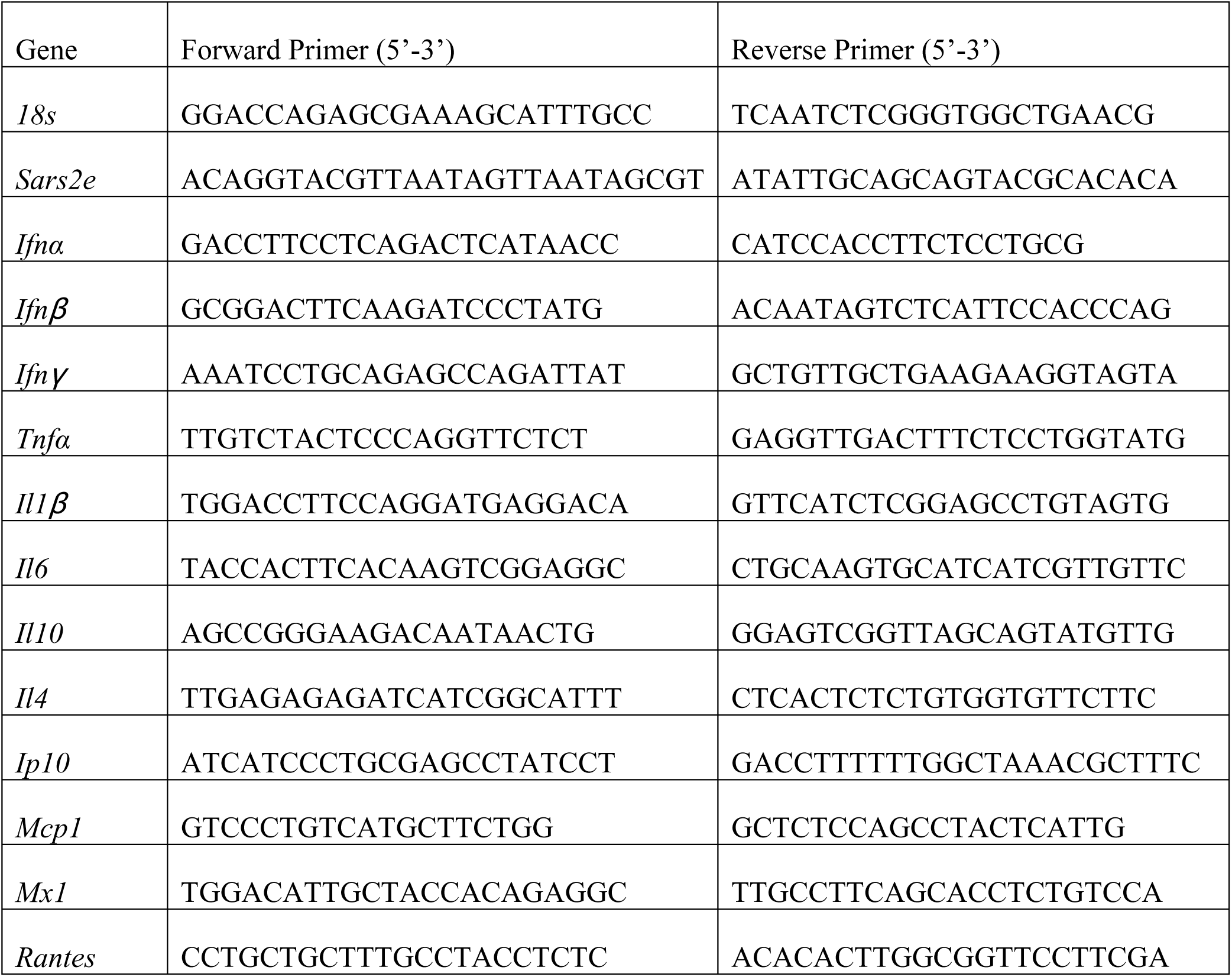
List of RT-qPCR primers.

20 ng cDNA was amplified for each replicate, with each animal specimen being assayed in duplicate for each gene, using an iTaq Universal SYBR Green supermix reagent kit (BioRad, 1725124, Hercules, California, USA), in a CFX96 thermocycler (BioRad, Hercules, California, USA). The cycling parameters for PCR for 40 cycles were as follows: initial polymerase activation at 95℃ for 30 s, denaturation at 95℃ for 10 s, and annealing/extension with plate read at 60℃ for 30 s. The relative fold gene expression for each sample was calculated based on the Livak delta-delta Ct method(64).

### Histopathology and immunostaining

Formalin-fixed whole skull sections in 70% ethanol from the above-described necropsy were subsequently paraffin-embedded and then sectioned at 5 µm thickness along the sagittal plane. Histopathological evaluation was performed on deparaffinized sections stained by routine hematoxylin-and-eosin (H&E) staining. Testing for the SARS-CoV-2 S viral antigen was performed using a standard colorimetric indirect horseradish peroxidase (HRP) IHC protocol modified from a previously described protocol(^37, 38^) using a rabbit anti-SARS-CoV-2 S protein antibody (Abcam, ab272504, Cambridge, UK) at 1:5000 dilution (0.2 µg/mL). Heat-mediated antigen retrieval at pH 6 using citrate buffer was performed. Specifically, the primary antibody was detected using the ImmPRESS® HRP Horse Anti-Rabbit IgG PLUS Polymer Kit (Vector Laboratories, MP-7801-15, Newark, California, USA) following manufacturer instructions.

Counterstaining was achieved with Mayer’s hematoxylin (Sigma-Aldrich, MHS16-500 mL, St. Louis, Missouri, USA). For two-color IF staining, the above IHC protocol was modified such that the anti-SARS-CoV-2 S primary antibody was at 1:1000 dilution (1 µg/mL) in background-reducing antibody diluent (Dako, S302283-2, Santa Clara, California, USA). A second primary antibody to detect a different cell marker antigen was simultaneously used with the SARS-CoV-2 S primary antibody at the following dilutions: TUBB3/Tuj1 (GeneTex, GTX85469, Irvine, California, USA, 1:100), GFAP (GeneTex, GTX85454, Irvine, California, USA, 1:500), and ACE2 (R&D Systems, AF933, Minneapolis, MN, USA, 1:250); IBA1 (GeneTex, GTX637629, Irvine, California, USA, 1:100) was used with a different mouse SARS-CoV-2 S antibody (GeneTex, GTX632604, Irvine, California, USA, 1:1000). The primary antibodies were then visualized using secondary antibodies conjugated with the appropriate listed fluorophores: goat anti-rabbit IgG Alexa Fluor 568 (Invitrogen, A-11011, Waltham, Massachusetts, USA, 1:1000), goat anti-chicken IgY Alexa Fluor 488 (Invitrogen, A-11039, Waltham, Massachusetts, USA, 1:2000), and goat anti-mouse IgG Alexa Fluor 555 (Invitrogen, A-32727, Waltham, Massachusetts, USA, 1:2000).

### Graph creation and statistical analysis

Statistical analysis was performed, and graphs were created in GraphPad Prism 10.2.3.

## ACKNOWLEDGEMENTS

We thank Dr. Kempaiah Rayavera Kempaiah for providing the initial samples. We also thank Dr. Vivian Tat for her help in providing technical assistance. We appreciate Dr. Ping Wu’s technical advice in neurobiology.

This work was supported by grant from the National Institute of Allergy and Infectious Diseases (HHSN272201700040I to DB and CKT).

---

**Supplemental Figure 1. Histopathological analysis of SARS-CoV-2-infected mice brains.** Micrograph of whole skull section (right hemisphere) of infected AC70 mouse at 5 dpi.show a lack of abnormalities or differences between the mock-infected sections (A-D) and sections from the SARS-2-infected brains (E-H). Magnification 10X.

**Supplemental Figure 2. Immunohistochemical analysis of SARS-CoV-2 antigen in the brain at 4 dpi.** Images of FFPE skull sections of infected AC70 mice at 4 dpi. (A) Whole left hemisphere, 10X; (B) pre-frontal cortex, 40X; (C) striatum, 40X; (D) thalamus, 40X; (E) hypothalamus, 40X; (F) hippocampus, 40X; (G) cerebellum, 40X; (H) medulla oblongata, 40X. Blue nuclei indicate hematoxylin counter-staining.

**Supplemental Figure 3. Immunohistochemical analysis of SARS-CoV-2 antigen in the brain at 5 dpi.** Micrograph of whole skull section (right hemisphere) of infected AC70 mouse at 5 dpi.

## REFERENCES

1. Organization WH. Pneumonia of unknown cause – China. World Health Organization-Disease Outbreak News-Item 2020.

2. Dong E, Du H, Gardner L. An interactive web-based dashboard to track COVID-19 in real time. The Lancet Infectious Diseases. 2020;20(5):533–4. doi: 10.1016/S1473-3099(20)30120-1.

3. Coronaviridae Study Group of the International Committee on Taxonomy of V. The species Severe acute respiratory syndrome-related coronavirus: classifying 2019-nCoV and naming it SARS-CoV-2. Nat Microbiol. 2020;5(4):536–44. Epub 2020/03/02. doi: 10.1038/s41564-020-0695-z. PubMed PMID: 32123347.

4. Baig AM, Sanders EC. Potential neuroinvasive pathways of SARS-CoV-2: Deciphering the spectrum of neurological deficit seen in coronavirus disease-2019 (COVID-19). J Med Virol. 2020;92(10):1845–57. Epub 2020/06/04. doi: 10.1002/jmv.26105. PubMed PMID: 32492193; PMCID: PMC7300748.

5. Bauer L, Laksono BM, de Vrij FMS, Kushner SA, Harschnitz O, van Riel D. The neuroinvasiveness, neurotropism, and neurovirulence of SARS-CoV-2. Trends in Neurosciences. 2022;45(5):358–68. doi: 10.1016/j.tins.2022.02.006.

6. DosSantos MF, Devalle S, Aran V, Capra D, Roque NR, Coelho-Aguiar JM, Spohr T, Subilhaga JG, Pereira CM, D’Andrea Meira I, Niemeyer Soares Filho P, Moura-Neto V. Neuromechanisms of SARS-CoV-2: A Review. Front Neuroanat. 2020;14(37):37. Epub 2020/07/03. doi: 10.3389/fnana.2020.00037. PubMed PMID: 32612515; PMCID: PMC7308495.

7. Huang YH, Jiang D, Huang JT. SARS-CoV-2 Detected in Cerebrospinal Fluid by PCR in a Case of COVID-19 Encephalitis. Brain Behav Immun. 2020;87:149. Epub 2020/05/11. doi: 10.1016/j.bbi.2020.05.012. PubMed PMID: 32387508; PMCID: PMC7202824.

8. Solomon IH, Normandin E, Bhattacharyya S, Mukerji SS, Keller K, Ali AS, Adams G, Hornick JL, Padera RF, Sabeti P. Neuropathological Features of Covid-19. New England Journal of Medicine. 2020;383(10):989–92. doi: 10.1056/NEJMc2019373.

9. Solomon T. Neurological infection with SARS-CoV-2 — the story so far. Nature Reviews Neurology. 2021;17(2):65–6. doi: 10.1038/s41582-020-00453-w.

10. Song E, Zhang C, Israelow B, Lu-Culligan A, Prado AV, Skriabine S, Lu P, Weizman O- E, Liu F, Dai Y, Szigeti-Buck K, Yasumoto Y, Wang G, Castaldi C, Heltke J, Ng E, Wheeler J, Alfajaro MM, Levavasseur E, Fontes B, Ravindra NG, Van Dijk D, Mane S, Gunel M, Ring A, Kazmi SAJ, Zhang K, Wilen CB, Horvath TL, Plu I, Haik S, Thomas J-L, Louvi A, Farhadian SF, Huttner A, Seilhean D, Renier N, Bilguvar K, Iwasaki A. Neuroinvasion of SARS-CoV-2 in human and mouse brainNeuroinvasion of SARS-CoV-2 in humans and mice. Journal of Experimental Medicine. 2021;218(3). doi: 10.1084/jem.20202135.

11. Arbour N, Day R, Newcombe J, Talbot PJ. Neuroinvasion by Human Respiratory Coronaviruses. Journal of Virology. 2000;74(19):8913–21. doi: doi:10.1128/JVI.74.19.8913-8921.2000.

12. Desforges M, Le Coupanec A, Brison É, Meessen-Pinard M, Talbot PJ, editors. Neuroinvasive and Neurotropic Human Respiratory Coronaviruses: Potential Neurovirulent Agents in Humans 2014; New Delhi: Springer India.

13. Al-Aly Z, Xie Y, Bowe B. High-dimensional characterization of post-acute sequelae of COVID-19. Nature. 2021;594(7862):259-64. doi: 10.1038/s41586-021-03553-9.

14. Proal AD, VanElzakker MB. Long COVID or Post-acute Sequelae of COVID-19 (PASC): An Overview of Biological Factors That May Contribute to Persistent Symptoms. Frontiers in Microbiology. 2021;12. doi: 10.3389/fmicb.2021.698169.

15. Jeong GU, Kwon H-J, Ng WH, Liu X, Moon HW, Yoon GY, Shin HJ, Lee I-C, Ling ZL, Spiteri AG, King NJC, Taylor A, Chae JS, Kim C, Ahn D-G, Kim K-D, Ryu YB, Kim S-J, Mahalingam S, Kwon Y-C. Ocular tropism of SARS-CoV-2 in animal models with retinal inflammation via neuronal invasion following intranasal inoculation. Nature Communications. 2022;13(1):7675. doi: 10.1038/s41467-022-35225-1.

16. Morris M, Zohrabian VM. Neuroradiologists, Be Mindful of the Neuroinvasive Potential of COVID-19. American Journal of Neuroradiology. 2020. doi: 10.3174/ajnr.A6551.

17. Natoli S, Oliveira V, Calabresi P, Maia LF, Pisani A. Does SARS-Cov-2 invade the brain? Translational lessons from animal models. Eur J Neurol. 2020;27(9):1764–73. Epub 2020/04/26. doi: 10.1111/ene.14277. PubMed PMID: 32333487; PMCID: PMC7267377.

18. Paniz-Mondolfi A, Bryce C, Grimes Z, Gordon RE, Reidy J, Lednicky J, Sordillo EM, Fowkes M. Central nervous system involvement by severe acute respiratory syndrome coronavirus-2 (SARS-CoV-2). J Med Virol. 2020;92(7):699–702. Epub 2020/04/22. doi: 10.1002/jmv.25915. PubMed PMID: 32314810; PMCID: PMC7264598.

19. Meinhardt J, Radke J, Dittmayer C, Franz J, Thomas C, Mothes R, Laue M, Schneider J, Brunink S, Greuel S, Lehmann M, Hassan O, Aschman T, Schumann E, Chua RL, Conrad C, Eils R, Stenzel W, Windgassen M, Rossler L, Goebel HH, Gelderblom HR, Martin H, Nitsche A, Schulz-Schaeffer WJ, Hakroush S, Winkler MS, Tampe B, Scheibe F, Kortvelyessy P, Reinhold D, Siegmund B, Kuhl AA, Elezkurtaj S, Horst D, Oesterhelweg L, Tsokos M, Ingold-Heppner B, Stadelmann C, Drosten C, Corman VM, Radbruch H, Heppner FL. Olfactory transmucosal SARS-CoV-2 invasion as a port of central nervous system entry in individuals with COVID-19. Nat Neurosci. 2021;24(2):168–75. Epub 2020/12/02. doi: 10.1038/s41593-020-00758-5. PubMed PMID: 33257876.

20. Jeong GU, Lyu J, Kim K-D, Chung YC, Yoon GY, Lee S, Hwang I, Shin W-H, Ko J, Lee J-Y, Kwon Y-C. SARS-CoV-2 Infection of Microglia Elicits Proinflammatory Activation and Apoptotic Cell Death. Microbiology Spectrum. 2022;10(3):e01091–22. doi: doi:10.1128/spectrum.01091-22.

21. Hoffmann M, Kleine-Weber H, Pöhlmann S. A Multibasic Cleavage Site in the Spike Protein of SARS-CoV-2 Is Essential for Infection of Human Lung Cells. Molecular Cell. 2020;78(4):779–84.e5. doi: 10.1016/j.molcel.2020.04.022.

22. Hoffmann M, Kleine-Weber H, Schroeder S, Kruger N, Herrler T, Erichsen S, Schiergens TS, Herrler G, Wu NH, Nitsche A, Muller MA, Drosten C, Pohlmann S. SARS-CoV-2 Cell Entry Depends on ACE2 and TMPRSS2 and Is Blocked by a Clinically Proven Protease Inhibitor. Cell. 2020;181(2):271–80 e8. Epub 2020/03/07. doi: 10.1016/j.cell.2020.02.052. PubMed PMID: 32142651; PMCID: PMC7102627.

23. Chen R, Wang K, Yu J, Howard D, French L, Chen Z, Wen C, Xu Z. The Spatial and Cell-Type Distribution of SARS-CoV-2 Receptor ACE2 in the Human and Mouse Brains. Frontiers in Neurology. 2021;11(1860). doi: 10.3389/fneur.2020.573095.

24. Theil D, Derfuss T, Paripovic I, Herberger S, Meinl E, Schueler O, Strupp M, Arbusow V, Brandt T. Latent Herpesvirus Infection in Human Trigeminal Ganglia Causes Chronic Immune Response. The American Journal of Pathology. 2003;163(6):2179–84. doi: 10.1016/S0002-9440(10)63575-4.

25. Beier KT. The Serendipity of Viral Trans-Neuronal Specificity: More Than Meets the Eye. Front Cell Neurosci. 2021;15:720807. Epub 2021/10/22. doi: 10.3389/fncel.2021.720807. PubMed PMID: 34671244; PMCID: PMC8521040.

26. Shimeld C, Efstathiou S, Hill T. Tracking the Spread of a *lacZ*-Tagged Herpes Simplex Virus Type 1 between the Eye and the Nervous System of the Mouse: Comparison of Primary and Recurrent Infection. Journal of Virology. 2001;75(11):5252–62. doi: doi:10.1128/jvi.75.11.5252-5262.2001.

27. Ptaszyńska-Sarosiek I, Dunaj J, Zajkowska A, Niemcunowicz-Janica A, Król M, Pancewicz S, Zajkowska J. Post-mortem detection of six human herpesviruses (HSV-1, HSV-2, VZV, EBV, CMV, HHV-6) in trigeminal and facial nerve ganglia by PCR. PeerJ. 2019;6:e6095. doi: 10.7717/peerj.6095.

28. Kamel HAM, Toland J. Trigeminal Nerve Anatomy. American Journal of Roentgenology. 2001;176(1):247–51. doi: 10.2214/ajr.176.1.1760247. PubMed PMID: 11133576.

29. Schaefer ML, Böttger B, Silver WL, Finger TE. Trigeminal collaterals in the nasal epithelium and olfactory bulb: A potential route for direct modulation of olfactory information by trigeminal stimuli. Journal of Comparative Neurology. 2002;444(3):221–6. doi: 10.1002/cne.10143.

30. Tremblay C, Frasnelli J. Olfactory and Trigeminal Systems Interact in the Periphery. Chemical Senses. 2018;43(8):611–6. doi: 10.1093/chemse/bjy049.

31. Romano N, Federici M, Castaldi A. Imaging of cranial nerves: a pictorial overview. Insights into Imaging. 2019;10(1):33. doi: 10.1186/s13244-019-0719-5.

32. Fagre A, Lewis J, Eckley M, Zhan S, Rocha SM, Sexton NR, Burke B, Geiss B, Peersen O, Bass T, Kading R, Rovnak J, Ebel GD, Tjalkens RB, Aboellail T, Schountz T. SARS-CoV-2 infection, neuropathogenesis and transmission among deer mice: Implications for spillback to New World rodents. PLOS Pathogens. 2021;17(5):e1009585. doi: 10.1371/journal.ppat.1009585.

33. Zhang AJ, Lee AC, Chu H, Chan JF, Fan Z, Li C, Liu F, Chen Y, Yuan S, Poon VK, Chan CC, Cai JP, Wu KL, Sridhar S, Chan YS, Yuen KY. SARS-CoV-2 infects and damages the mature and immature olfactory sensory neurons of hamsters. Clin Infect Dis. 2020. Epub 2020/07/16. doi: 10.1093/cid/ciaa995. PubMed PMID: 32667973; PMCID: PMC7454453.

34. Pekny M, Wilhelmsson U, Pekna M. The dual role of astrocyte activation and reactive gliosis. Neuroscience Letters. 2014;565:30–8. doi: 10.1016/j.neulet.2013.12.071.

35. Pekny M, Pekna M. Astrocyte Reactivity and Reactive Astrogliosis: Costs and Benefits. Physiological Reviews. 2014;94(4):1077–98. doi: 10.1152/physrev.00041.2013. PubMed PMID: 25287860.

36. Garman RH. Histology of the Central Nervous System. Toxicologic Pathology. 2011;39(1):22–35. doi: 10.1177/0192623310389621. PubMed PMID: 21119051.

37. Tseng C-TK, Huang C, Newman P, Wang N, Narayanan K, Watts DM, Makino S, Packard MM, Zaki SR, Chan T-s, Peters CJ. Severe Acute Respiratory Syndrome Coronavirus Infection of Mice Transgenic for the Human Angiotensin-Converting Enzyme 2 Virus Receptor. Journal of Virology. 2007;81(3):1162–73. doi: doi:10.1128/jvi.01702-06.

38. Yoshikawa N, Yoshikawa T, Hill T, Huang C, Watts DM, Makino S, Milligan G, Chan T, Peters CJ, Tseng C-TK. Differential Virological and Immunological Outcome of Severe Acute Respiratory Syndrome Coronavirus Infection in Susceptible and Resistant Transgenic Mice Expressing Human Angiotensin-Converting Enzyme 2. Journal of Virology. 2009;83(11):5451–65. doi: doi:10.1128/JVI.02272-08.

39. Ogando NS, Dalebout TJ, Zevenhoven-Dobbe JC, Limpens RWAL, van der Meer Y, Caly L, Druce J, de Vries JJC, Kikkert M, Bárcena M, Sidorov I, Snijder EJ. SARS-coronavirus-2 replication in Vero E6 cells: replication kinetics, rapid adaptation and cytopathology. Journal of General Virology. 2020;101(9):925–40. doi: 10.1099/jgv.0.001453.

40. Bulfamante G, Bocci T, Falleni M, Campiglio L, Coppola S, Tosi D, Chiumello D, Priori A. Brainstem neuropathology in two cases of COVID-19: SARS-CoV-2 trafficking between brain and lung. Journal of Neurology. 2021;268(12):4486–91. doi: 10.1007/s00415-021-10604-8.

41. Bulfamante G CD, Canevini MP, Priori A, Mazzanti M, Centanni S, et al. First ultrastructural autoptic findings of SARS-Cov-2 in olfactory pathways and brainstem.. Minerva Anestesiol 2020;86:678–9. doi: 10.23736/S0375-9393.20.14772-2.

42. Emmi A, Rizzo S, Barzon L, Sandre M, Carturan E, Sinigaglia A, Riccetti S, Della Barbera M, Boscolo-Berto R, Cocco P, Macchi V, Antonini A, De Gaspari M, Basso C, De Caro R, Porzionato A. Detection of SARS-CoV-2 viral proteins and genomic sequences in human brainstem nuclei. npj Parkinson’s Disease. 2023;9(1):25. doi: 10.1038/s41531-023-00467-3.

43. Viszlayová D, Sojka M, Dobrodenková S, Szabó S, Bilec O, Turzová M, Ďurina J, Baloghová B, Borbély Z, Kršák M. SARS-CoV-2 RNA in the Cerebrospinal Fluid of a Patient with Long COVID. Therapeutic Advances in Infectious Disease. 2021;8:20499361211048572. doi: 10.1177/20499361211048572. PubMed PMID: 34659752.

44. Kemp WJ, Tubbs RS, Cohen-Gadol AA. The Innervation of the Cranial Dura Mater: Neurosurgical Case Correlates and a Review of the Literature. World Neurosurgery. 2012;78(5):505–10. doi: 10.1016/j.wneu.2011.10.045.

45. Pulous FE, Cruz-Hernández JC, Yang C, Kaya Z, Paccalet A, Wojtkiewicz G, Capen D, Brown D, Wu JW, Schloss MJ, Vinegoni C, Richter D, Yamazoe M, Hulsmans M, Momin N, Grune J, Rohde D, McAlpine CS, Panizzi P, Weissleder R, Kim D-E, Swirski FK, Lin CP, Moskowitz MA, Nahrendorf M. Cerebrospinal fluid can exit into the skull bone marrow and instruct cranial hematopoiesis in mice with bacterial meningitis. Nature Neuroscience. 2022;25(5):567–76. doi: 10.1038/s41593-022-01060-2.

46. Brann DH, Tsukahara T, Weinreb C, Lipovsek M, Van den Berge K, Gong B, Chance R, Macaulay IC, Chou HJ, Fletcher RB, Das D, Street K, de Bezieux HR, Choi YG, Risso D, Dudoit S, Purdom E, Mill J, Hachem RA, Matsunami H, Logan DW, Goldstein BJ, Grubb MS, Ngai J, Datta SR. Non-neuronal expression of SARS-CoV-2 entry genes in the olfactory system suggests mechanisms underlying COVID-19-associated anosmia. Sci Adv. 2020;6(31):eabc5801. Epub 2020/09/17. doi: 10.1126/sciadv.abc5801. PubMed PMID: 32937591.

47. Hamming I, Timens W, Bulthuis ML, Lely AT, Navis G, van Goor H. Tissue distribution of ACE2 protein, the functional receptor for SARS coronavirus. A first step in understanding SARS pathogenesis. J Pathol. 2004;203(2):631–7. Epub 2004/05/14. doi: 10.1002/path.1570. PubMed PMID: 15141377; PMCID: PMC7167720.

48. Pellegrini L, Albecka A, Mallery DL, Kellner MJ, Paul D, Carter AP, James LC, Lancaster MA. SARS-CoV-2 Infects the Brain Choroid Plexus and Disrupts the Blood-CSF Barrier in Human Brain Organoids. Cell Stem Cell. 2020;27(6):951–61.e5. doi: 10.1016/j.stem.2020.10.001.

49. Borrow P, Evans CF, Oldstone MB. Virus-induced immunosuppression: immune system-mediated destruction of virus-infected dendritic cells results in generalized immune suppression. Journal of Virology. 1995;69(2):1059–70. doi: doi:10.1128/jvi.69.2.1059-1070.1995.

50. Libbey JE FR. Virus-induced Immunosuppression. In: In: Brogden KA GJ, editors., editor. Polymicrobial Diseases Washington (DC): ASM Press; 2002.

51. McChesney MB, Fujinami RS, Lerche NW, Marx PA, Oldstone MBA. Virus-Induced Immunosuppression: Infection of Peripheral Blood Mononuclear Cells and Suppression of Immunoglobulin Synthesis During Natural Measles Virus Infection of Rhesus Monkeys. The Journal of Infectious Diseases. 1989;159(4):757–60. doi: 10.1093/infdis/159.4.757.

52. Hollville E, Romero SE, Deshmukh M. Apoptotic cell death regulation in neurons. The FEBS Journal. 2019;286(17):3276–98. doi: 10.1111/febs.14970.

53. Yakovlev AG, Faden AI. Mechanisms of neural cell death: Implications for development of neuroprotective treatment strategies. NeuroRX. 2004;1(1):5–16. doi: 10.1602/neurorx.1.1.5.

54. Kole AJ, Annis RP, Deshmukh M. Mature neurons: equipped for survival. Cell Death & Disease. 2013;4(6):e689-e. doi: 10.1038/cddis.2013.220.

55. Daubner SC, Le T, Wang S. Tyrosine hydroxylase and regulation of dopamine synthesis. Archives of Biochemistry and Biophysics. 2011;508(1):1–12. doi: 10.1016/j.abb.2010.12.017.

56. Zhuang Y, Krumm B, Zhang H, Zhou XE, Wang Y, Huang X-P, Liu Y, Cheng X, Jiang Y, Jiang H, Zhang C, Yi W, Roth BL, Zhang Y, Xu HE. Mechanism of dopamine binding and allosteric modulation of the human D1 dopamine receptor. Cell Research. 2021;31(5):593–6. doi: 10.1038/s41422-021-00482-0.

57. Kim HJ, Tsao JW, Stanfill AG. The current state of biomarkers of mild traumatic brain injury. JCI Insight. 2018;3(1). doi: 10.1172/jci.insight.97105.

58. Graham NSN, Zimmerman KA, Moro F, Heslegrave A, Maillard SA, Bernini A, Miroz J- P, Donat CK, Lopez MY, Bourke N, Jolly AE, Mallas E-J, Soreq E, Wilson MH, Fatania G, Roi D, Patel MC, Garbero E, Nattino G, Baciu C, Fainardi E, Chieregato A, Gradisek P, Magnoni S, Oddo M, Zetterberg H, Bertolini G, Sharp DJ. Axonal marker neurofilament light predicts long-term outcomes and progressive neurodegeneration after traumatic brain injury. Science Translational Medicine. 2021;13(613):eabg9922. doi: doi:10.1126/scitranslmed.abg9922.

59. Shahim P, Gren M, Liman V, Andreasson U, Norgren N, Tegner Y, Mattsson N, Andreasen N, Öst M, Zetterberg H, Nellgård B, Blennow K. Serum neurofilament light protein predicts clinical outcome in traumatic brain injury. Scientific Reports. 2016;6(1):36791. doi: 10.1038/srep36791.

60. Lee D, Cho Y, Ko Y, Heo NH, Kang HG, Han S. Neuron-specific enolase level as a predictor of neurological outcome in near-hanging patients: A retrospective multicenter study. PLOS ONE. 2021;16(2):e0246898. doi: 10.1371/journal.pone.0246898.

61. Rech TH, Vieira SR, Nagel F, Brauner JS, Scalco R. Serum neuron-specific enolase as early predictor of outcome after in-hospital cardiac arrest: a cohort study. Critical Care. 2006;10(5):R133. doi: 10.1186/cc5046.

62. Yap TF, Hsu JC, Liu Z, Rayavara K, Tat V, Chien-Te KT, Preston DJ. Efficacy and Self-Similarity of SARS-CoV-2 Thermal Decontamination. Journal of Hazardous Materials. 2021:127709. doi: 10.1016/j.jhazmat.2021.127709.

63. Reed Lj, Muench H. A SIMPLE METHOD OF ESTIMATING FIFTY PER CENT ENDPOINTS12. American Journal of Epidemiology. 1938;27(3):493–7. doi: 10.1093/oxfordjournals.aje.a118408.

64. Livak KJ, Schmittgen TD. Analysis of Relative Gene Expression Data Using Real-Time Quantitative PCR and the 2−ΔΔCT Method. Methods. 2001;25(4):402–8. doi: 10.1006/meth.2001.1262.

